# FOXA1 directs H3K4 monomethylation at enhancers via recruitment of the methyltransferase MLL3

**DOI:** 10.1101/069450

**Authors:** Kamila M. Jozwik, Igor Chernukhin, Aurelien A. Serandour, Sankari Nagarajan, Jason S. Carroll

**Affiliations:** Cancer Research UK Cambridge Institute, University of Cambridge, Robinson Way, Cambridge, UK, CB2 ORE; European Molecular Biology Laboratory, Genome Biology Unit, 69117, Heidelberg, Germany

**Keywords:** breast cancer, enhancers, H3K4me1, FOXA1, MLL3

## Abstract

FOXA1 is a pioneer factor that is important in hormone dependent cancer cells to stabilise nuclear receptors, such as estrogen receptor (ER) to chromatin. FOXA1 binds to enhancers regions that are enriched in H3K4mono- and dimethylation (H3K4me1, H3K4me2) histone marks and evidence suggests that these marks are requisite events for FOXA1 to associate with enhancers to initate subsequent gene expression events. However, exogenous expression of FOXA1 has been shown to induce H3K4me1 and H3K4me2 signal at enhancer elements and the order of events and the functional importance of these events is not clear. We performed a FOXA1 Rapid Immunoprecipitation Mass Spectrometry of Endogenous Proteins (RIME) screen in ERα-positive MCF-7 breast cancer cells in order to identify FOXA1 interacting partners and we found histone-lysine N-methyltransferase (MLL3) as the top FOXA1 interacting protein. MLL3 is typically thought to induce H3K4me3 at promoter regions, but recent findings suggest it may contribute to H3K4me1 deposition, in line with our observation that MLL3 associates with an enhancer specific protein. We performed MLL3 ChIP-seq in breast cancer cells and unexpectedly found that MLL3 binds mostly at non-promoter regions enhancers, in contrast to the prevailing hypothesis. MLL3 was shown to occupy regions marked by FOXA1 occupancy and as expected, H3K4me1 and H3K4me2. MLL3 binding was dependent on FOXA1, indicating that FOXA1 recruits MLL3 to chromatin. Motif analysis and subsequent genomic mapping revealed a role for Grainy head like protein-2 (GRHL2) which was shown to co-occupy regions of the chromatin with MLL3. Regions occupied by all three factors, namely FOXA1, MLL3 and GRHL2, were most enriched in H3K4me1. MLL3 silencing decreased H3K4me1 at enhancer elements, but had no appreciable impact on H3K4me3 at enhancer elements. We identify a complex relationship between FOXA1, MLL3 and H3K4me1 at enhancers in breast cancer and propose a mechanism whereby the pioneer factor FOXA1 can interact with a chromatin modifier MLL3, recruiting it to chromatin to facilitate the deposition of H3K4me1 histone marks, subsequently demarcating active enhancer elements.

## INTRODUCTION

FOXA1 (Forkhead Box Protein A1) is a pioneer factor (Jozwik and Carroll, 2012), which binds to condensed chromatin and allows subsequent binding of other transcription factors. FOXA1 contributes to chromatin opening to facilitate binding of Estrogen Receptor α (ER) in breast cancer (Carroll et al., 2005) and Androgen Receptor (AR) in prostate and breast cancer cells (Robinson et al., 2011; Sahu et al., 2011; Yang and Yu, 2015). ER is a driver of cell proliferation and tumour growth and ER positive breast cancer accounts for over 70 per cent of all breast cancers (Curtis et al., 2012). Recent evidence has shown that FOXA1 is essential for almost all ER binding events in breast cancer (Hurtado et al., 2011) and for ER functionality, yet our understanding of FOXA1 activity and the events involved in determining FOXA1-chromatin interactions are not well understood.

FOXA1 binding occurs at enhancer regions enriched in histone 3 lysine 4 mono and dimethylation (H3K4me1/me2) (Lupien et al., 2008). While it has been reported that FOXA1 binding requires H3K4me1/me2 marks (Lupien et al., 2008), more recent findings showed that exogenous expression of FOXA1 in the FOXA1-negative MDA-MB-231 cell line results in the acquisition of H3K4me1/me2 at FOXA1 bound sites (Serandour et al., 2011) suggesting that FOXA1 may actually contribute to deposition of the H3K4me1 and H3K4me2 marks, rather than associating with enhancers that are demarcated by the presence of these marks. Clearly, the order of these events is not resolved, yet the consequences of FOXA1 binding and H3K4me1/me2 signal result in a functional enhancer element that can recruit additional factors (such as ER) to drive expression of genes, including those involved in cell cycle progression.

Unlike H3K4me1 and H3K4me2 which is typically found at enhancer elements, H3K4me3 is typically observed at promoter regions and several investigations have associated the histone-lysine N-methyltransferase enzyme MLL3 with the deposition of H3K4me3 marks at promoters (Ardehali et al., 2011; Vermeulen and Timmers, 2010). More recently the MLL3/MLL4 complex has been implicated in regulation of H3K4me1 in mouse (Herz et al., 2012). Importantly MLL3 is mutated in a number of solid cancers, including 8-11% of breast cancers (Ellis et al., 2012; Wang et al., 2011), although a role for MLL3 in breast cancer and the functional consequences of these mutational events are not known. Silencing of MLL3 (and the related protein MLL2) has been shown to decrease the estrogen-mediated activation of HOXC6 in human placental choriocarcinoma (JAR cell line) and knockdown of either ERα or ERβ abolished estrogen-dependent recruitment of MLL2 and MLL3 onto the HOXC6 promoter in the JAR cell line. (Ansari et al., 2011).

We sought to discover novel proteins that interact with FOXA1 in ER-positive (ER+) breast cancer cells, by performing FOXA1 RIME (Rapid Immunoprecipitation-Mass Spectrometry of Endogenous proteins), an unbiased proteomic method that permits discovery of novel protein networks. This revealed a role for MLL3 as a critical chromatin regulatory protein at enhancer elements and as a factor that contributes to H3K4me1 deposition at these enhancers.

## RESULTS

### RIME analysis of FOXA1-associated proteins reveals interactions with MLL3 in breast cancer cells

We performed Rapid Immunoprecipitation Mass spectrometry of Endogenous protein (RIME) (Mohammed et al., 2013) of FOXA1 in MCF-7 breast cancer cells to identify endogenous FOXA1 interactors. Asychronous MCF-7 cells were grown in full media and five replicates of FOXA1 RIME were conducted. MLL3 was identified as one of the strongest and most reproducible interactors (Figure 1A) and the peptide coverage, number of unique peptides identified and Mascot score of MLL3 and FOXA1 are shown in Figure 1B. A full list of FOXA1 Interacting proteins is provided in Supplementary Table 1. We hypothesized that this interaction may be functional, as the pioneer factor FOXA1 binds at enhancer regions enriched in H3K4me1/me2 histone marks and FOXA1 has been shown to contribute to the acquisition of the H3K4me1/me2 mark (Sérandour et al., 2011). In addition, loss of MLL3 has been previously shown to correlate with reduced H3K4me1 at specific regions within the genome in mice (Herz et al., 2012). Our findings that MLL3 and FOXA1 physically interact in breast cancer cells, implies that FOXA1 may be able to directly recruit the enzyme that can add methyl groups to the H3K4 residue (Figure 1C).

**Figure 1.**
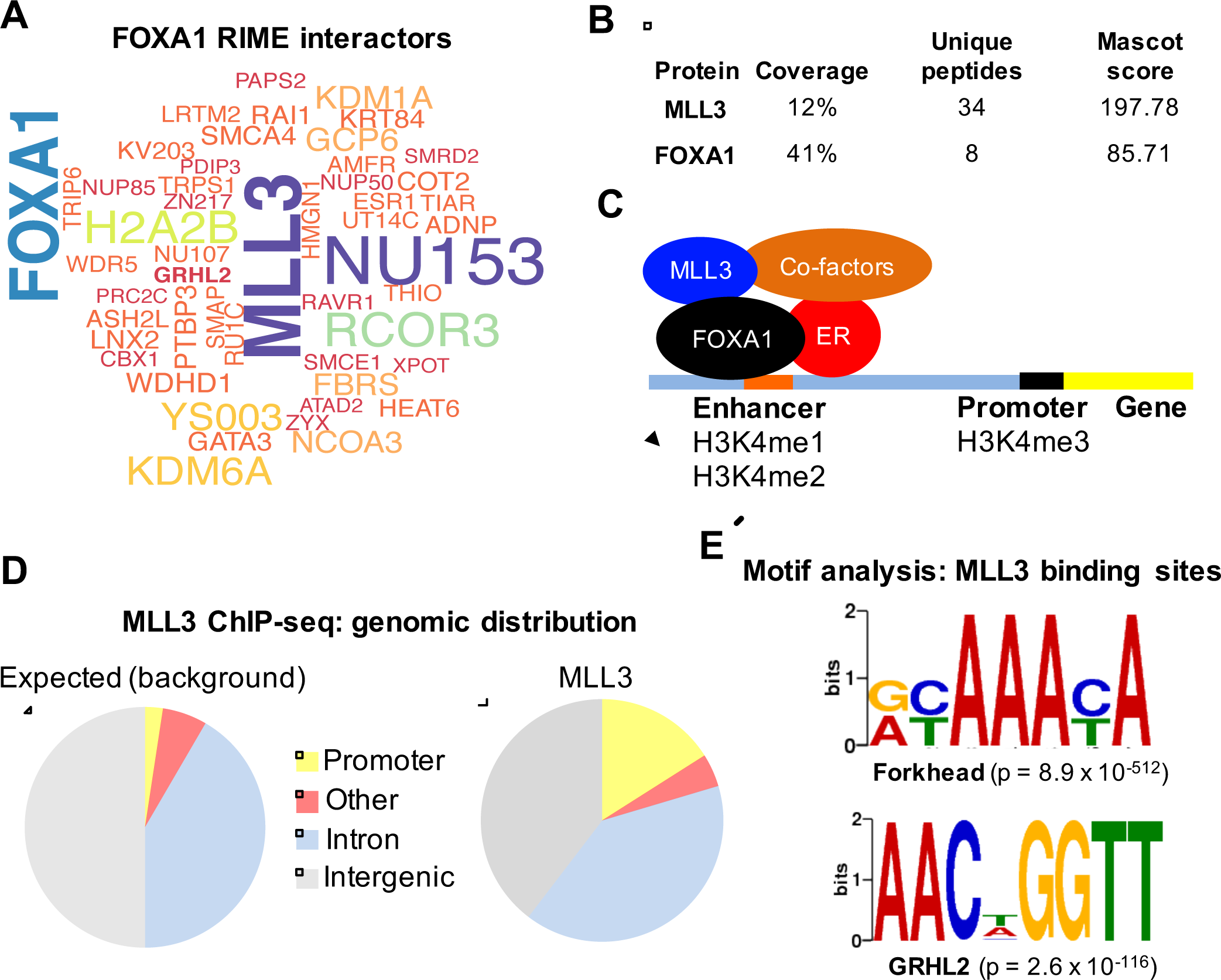
Purification of FOXA1-Associated Proteins Using RIME and Mapping of MLL3 binding genome-wide. **A.** FOXA1 interactome was discovered by performing RIME in MCF-7 breast cancer cells. The data is represented as a Wordcloud, where the size of protein names represent the strength/confidence of interactions based on the Mascot score. MLL3 was identified as one the strongest and most reproducible FOXA1 interacting proteins. **B.** Peptide coverage, number of unique peptides identified and Mascot score of MLL3 and FOXA1 following FOXA1 purification. **C.** Hypothesized mechanism of FOXA1 and MLL3 function. Our findings that MLL3 and FOXA1 physically interact in breast cancer cells implies that FOXA 1 could recruit the enzyme that can add methyl groups to histone 3 lysine 4. FOXA1 bound enhancers are demarcated by H3K4me1 and H3K4me2. **D.** MLL3 ChIP-seq was conducted and the genomic distribution of MLL3 peaks is shown, relative to the whole genome (the expected control values). Regions bound by MLL3 occurred mostly at enhancers, rather than promoters. **E.** *De novo* motif analysis of MLL3 binding sites. Motif analysis revealed an enrichment in Forkhead motif, the canonical motif bound by FOXA1 and motifs for the transcription factor grainyhead-like 2 protein (GRHL2).

### Global mapping of MLL3 binding sites shows enrichment at enhancers

Due to the size of MLL3 (~540kDa) and the fact that FOXA1 is the same size at the antibody heavy chain, we were unable to conduct Co-IP validation experiments of the newly discovered MLL3-FOXA1 interactions. However, we explored this putative interaction in several ways. We reversed the pulldown, purified MLL3 and could discover FOXA1 as an interacting protein by RIME (data not shown). In addition, we assessed the global interplay between MLL3 and FOXA1 binding events, by performed MLL3 and FOXA1 ChIP-seq. Asynchronous MCF-7 cells were grown and triplicate ChIP-seq experiments were conducted. MLL3 ChIP-seq was conducted using a custom antibody, that we validated by RIME and could show to be specific to MLL3 (Supplementary Figure 1). Data from all the replicates were pooled and peaks were called using MACS (Zhang et al., 2008) resulting in a total of 10,772 MLL3 binding events in MCF-7 cells. MLL3 has been previously implicated as an enzyme that contributes to H3K4me3 deposition at promoter regions (Ananthanarayanan et al., 2011), but our ChIP-seq data showed that MLL3 binding was mostly distributed at enhancer elements and intergenic regions (Figure 1D) with a smaller fraction distributed at promoters, similar to what has been observed for ER and FOXA1 (Carroll et al., 2005; Hurtado et al., 2011). Analysis of enriched DNA motifs within the MLL3 binding sites revealed an enrichment in Forkhead motifs, the canonical motif bound by FOXA1 (MEME evalue = 8.9e-512). In addition, motifs for the transcription factor grainyhead-like 2 protein (GRHL2) were identified (Figure 1E) (MEME evalue = 2.6e-116), although there is limited information linking GRHL2 and ER/FOXA1 signalling.

### MLL3 binding is dependent on FOXA1

ChIP-seq of FOXA1 and H3K4me1/me3 were performed in asynchronous MCF-7 cells in triplicate and peaks were called using MACS, revealing 23,375 FOXA1 peaks, 26,584 H3K4me1 and 13,478 H3K4me3 peaks. The binding of FOXA1 and H3K4me1/me3 was overlapped with the MLL3 binding sites. The majority (55.8%) of MLL3 binding events were co-bound by FOXA1 (Figure 2A) and since H3K4me1 is associated with FOXA1, it was not unexpected that MLL3/FOXA1 co-bound regions were also typically marked by H3K4me1 (Figure 2A). A small percentage (9.1%) of MLL3/FOXA1 co-bound regions were also marked by histone 3 lysine 4 trimethylation (H3K4me3) (Supplementary Figure 2). An example of a MLL3 and FOXA1 co-bound region, marked by both H3K4me1 and H3K4me3, is shown in Figure 2B. Heatmap visualisation of the FOXA1 binding and H3K4me1/me3 signal at the MLL3 binding events is shown in Figure 2C, indicating that a substantial degree of the MLL3 and FOXA1 co-bound regions also possess H3K4me1 signal.

**Figure 2.**
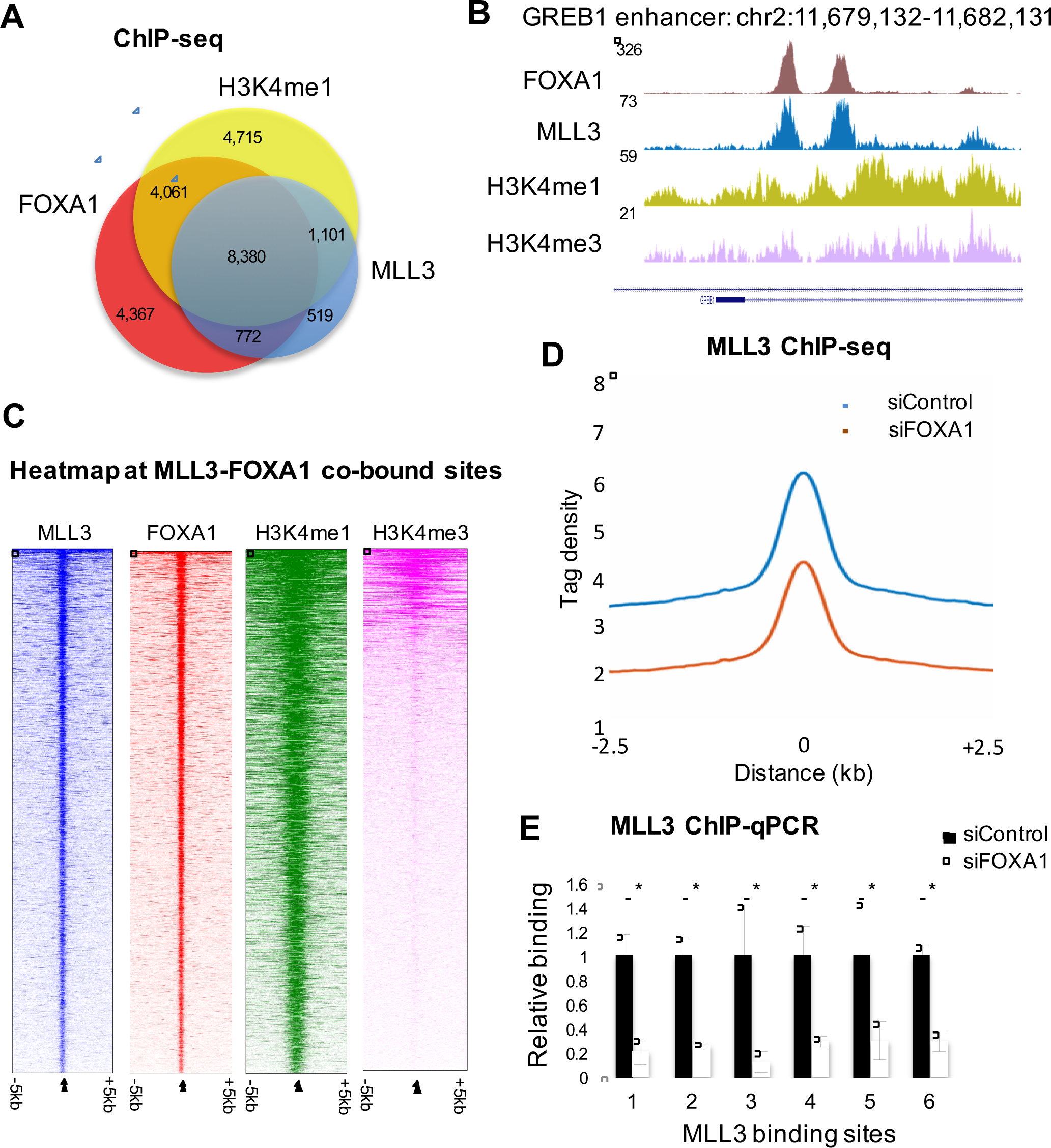
Co-binding of MLL3, FOXA1 and H3K4me1/me3 and mechanism of MLL3 recruitment. **A.** Overlap between MLL3, FOXA1 and H3K4me1 binding revealed by ChIP-seq. MLL3 binding sites were co-bound by FOXA1 and the histone marks. The numbers of peaks within each category is shown on the diagram. **B.** An example of MLL3, FOXA1 and H3K4me1/me3 co-bound region at the GREB1 enhancer. **C.** Heatmap of MLL3-FOXA1 co-bound regions showing binding signal intensity for FOXA1, MLL3, H3K4me1 and H3K4me3. Binding is ranked from the strongest to the weakest binding sites. **D.** Signal intensity plot representing changes in MLL3 ChIP-seq signal in siControl versus siFOXA1 tranfected conditions. Differentially bound sites needed to be detected in at least two replicates to be included. **E.** ChIP-qPCR validation of MLL3 binding sites that were lost upon FOXA1 silencing. The ChIP data is shown as relative change to siControl. * denotes p < 0.05, as determined by t-test.

Given that MLL3 was the top FOXA1 interacting protein (Figure 1A), that MLL3 binding sites were enriched for Forkhead motifs and that 55.8% of MLL3 binding events we also FOXA1 binding sites, we hypothesised that MLL3 was recruited to the chromatin by FOXA1. To assess this, MCF-7 cells were transfected with siControl or siRNA to FOXA1 and effective FOXA1 silencing was confirmed. Following FOXA1 silencing MLL3 ChIP-seq was conducted in triplicate independent biological replicates. MLL3 peaks were called in siControl or siFOXA1 transfected conditions. This resulted in a global decrease in MLL3 binding when FOXA1 was depleted (Figure 2D). The decreased MLL3 binding following silencing of FOXA1 was not due to a decrease in MLL3 expression, since MLL3 mRNA levels increased following FOXA1 silencing (Supplementary Figure 2). Six MLL3 binding sites were assessed using ChIP-qPCR, validating the dependence on FOXA1 for MLL3 binding to chromatin (Figure 2E). The importance of MLL3 for MCF-7 breast cancer cell growth was confirmed by silencing MLL3 (or FOXA1 as a positive control) and assessing the impact on cell confluence over time (Supplementary Figure 3).

### Chromatin properties at MLL3 binding events

As previously observed (Figure 1E), GRHL2 (grainyhead-like 2 protein) motifs were enriched within MLL3 binding events. GRHL2 was also found to be a FOXA1 interacting protein from the RIME experiments (Figure 1A), suggesting that the enrichment of GRHL2 motifs might represent a functional interaction between FOXA1 and GRHL2. The role of GRHL2 in breast cancer is currently unclear, with both pro-metastatic and anti-metastatic roles (Werner et al., 2013; Xiang et al., 2012). We performed GRHL2 ChIP-seq in MCF-7 cells in triplicate and GRHL2 peaks were called using MACS, revealing 30,143 GRHL2 binding sites. GRHL2 binding was overlaid with MLL3 and FOXA1 binding, revealing 5,585 regions that were occupied by all three factors (Figure 3A). An example of a co-occupied site is shown in Figure 3B. In total, 91.5% of MLL3 binding were co-occupied by FOXA1 and/or GRHL2. To gain insight into the mechanisms involved in the different cis-regulatory elements, we explored the seven different categories of binding, by investigating regions bound by a single factor (FOXA1 only, MLL3 only, GRHL2 only), two factors (FOXA1 and MLL3, MLL3 and GRHL2, GRHL2 and FOXA1) or all three factors and used them for further analyses. Only 8.5% of the MLL3 binding regions were not co-bound by FOXA1, GRHL2 or both, suggesting that MLL3 can not associate with chromatin without one of the associated transcription factors and the MLL3 only binding regions were subsequently eliminated from further analyses.

**Figure 3.**
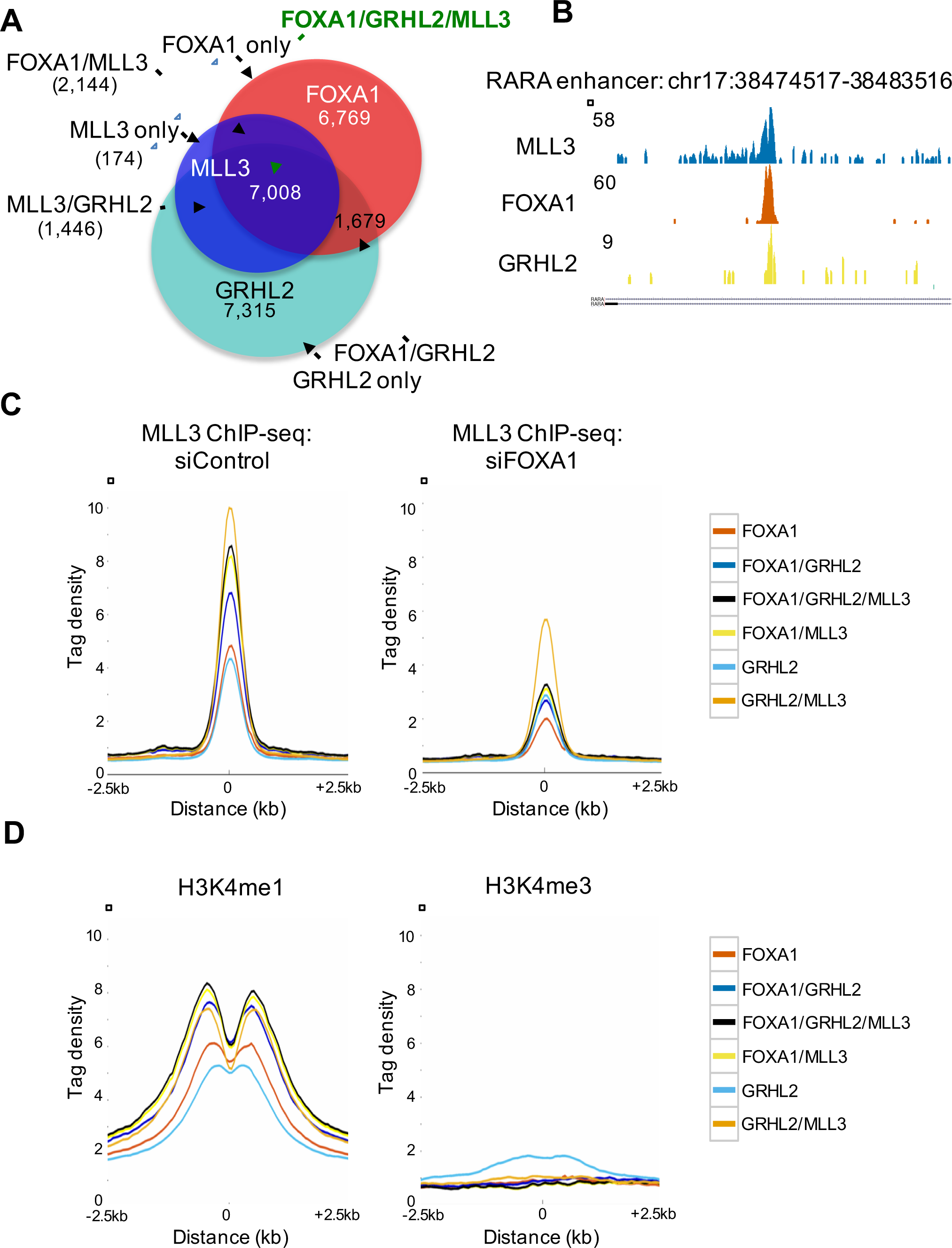
Functional distinction between regions bound by FOXA1, GRHL2 and MLL3. **A.** Venn diagram showing the overlap between MLL3, FOXA1 and GRHL2 binding regions, identifying the different categories of binding events. For subsequent analyses, we assessed the number of regions co-bound by one factor (FOXA1 only, MLL3 only, GRHL2 only), two factors (FOXA1 and MLL3, MLL3 and GRHL2, GRHL2 and FOXA1) and all three factors. **B.** An example of a MLL3, FOXA1 and GRHL2 co-bound region. **C.** Average MLL3 binding signal in siControl and siFOXA1 conditions at the different binding categories. Following FOXA1 silencing, MLL3 binding intensity was reduced at regions occupied by MLL3, FOXA1 and GRHL2, regions occupied by MLL3 and FOXA1 and to a lesser extent at regions occupied by MLL3 and GRHL2. **D.** H3K4me1/me3 distribution at the different binding regions. The most enriched H3K4me1 regions were those where MLL3 was recruited.

In control conditions, MLL3 binding was most enriched at sites co-occupied by either FOXA1, GRHL2 or both proteins together, suggesting that optimal MLL3-chromatin occupancy involves at least one of the additional transcription factors (Figure 3C). Following silencing of FOXA1, MLL3 binding was substantially reduced at two categories, the first was the regions bound by all three proteins and the second was the FOXA1 and MLL3 (but not GRHL2) regions. Interestingly, MLL3 binding signal at MLL3 and GRHL2 (but not FOXA1) occupied cis-regulatory elements were moderately affected by FOXA1 silencing, suggesting multiple modes of MLL3-chromatin occupancy (Figures 3C). This suggests that upon FOXA1 silencing, MLL3 binding sites were lost at any region where FOXA1 co-binds, even if GRHL2 is also present, but MLL3 binding is moderately affected at regions where GRHL2 is the sole associated protein with MLL3.

When the different MLL3 binding regions were integrated with the H3K4me1/me3 data, the most enriched regions were those where MLL3, FOXA1 and GRHL2 were co-bound and those where MLL3 and FOXA1 were co-bound, although any region occupied by MLL3 had increased H3K4me1 signal, relative to regions occupied by FOXA1 or GRHL2, but not MLL3 (Figure 3D). These findings indicate that the presence of MLL3 correlates with increased H3K4me1.

Since FOXA1 contributes to the establishment of enhancer elements that are subsequently used by transcription factors such as Estrogen Receptor (ER) in these breast cancer cells, we integrated the MLL3, FOXA1 and GRHL2 ChIP-seq data with ER binding information. As expected (Supplementary Figure 4), the regions bound by FOXA1 and MLL3 are commonly co-occupied by ER, in support of their role in establishing ER enhancer elements.

### H3K4me1 at enhancers elements is dependent on MLL3

Given that MLL3 binding was associated with regions enriched in H3K4me1 marks and MLL3 is a methyltransferase, we hypothesised that MLL3 contributes to the presence of this methyl mark at enhancer elements. To assess this, MCF-7 cells were transfected with siControl or siMLL3 and triplicate H3K4me1 and H3K4me3 ChIP-seq experiments were conducted and peaks were called using MACS. When MLL3 was silenced, deposition of H3K4me1 was substantially decreased at both enhancer elements and promoters (Figure 4A). We specifically assessed the changes in H3K4me1 at regions bound by both FOXA1 and MLL3, resulting in the identification of 776 FOXA1/MLL3 bound enhancers that had decreased H3K4me1 following MLL3 silencing (Figure 4B). There was no decreased H3K4me3 at either the enhancer elements or the promoter regions when MLL3 was silenced and a modest gain of signal at both promoters and enhancer elements (Figure 4C). We assessed the 776 enhancer elements with decreased H3K4me1 following MLL3 silencing for enriched pathways. GREAT revealed a number of enriched pathways, most of which were associated with transcriptional regulation (Figure 4D). Given the observation that regions bound by MLL3 possess the highest H3K4me1 signal (Figure 4E) and that H3K4me1 was depleted at enhancers when MLL3 was silenced, we postulate that H3K4me1 deposition is mediated by MLL3 at enhancer elements, as determined by FOXA1 and/or GRHL2 recruitment of MLL3. In support of this, MCF-7 cells were transfected with siControl or siFOXA1 and H3K4me1 ChIP-qPCR was conducted on a select number of loci (the genomic rgions and the relative factor binding is shown in Supplementary Figure 5. Following inhibition of FOXA1, H3K4me1 signal was dimished at some of the assessed loci (Supplementary Figure 6) and H3K27Ac was also decreased following FOXA1 inhibition (Supplementary Figure 7), imply that FOXA1 and MLL3 are required for transcriptional activity from the enhancer elements.

**Figure 4.**
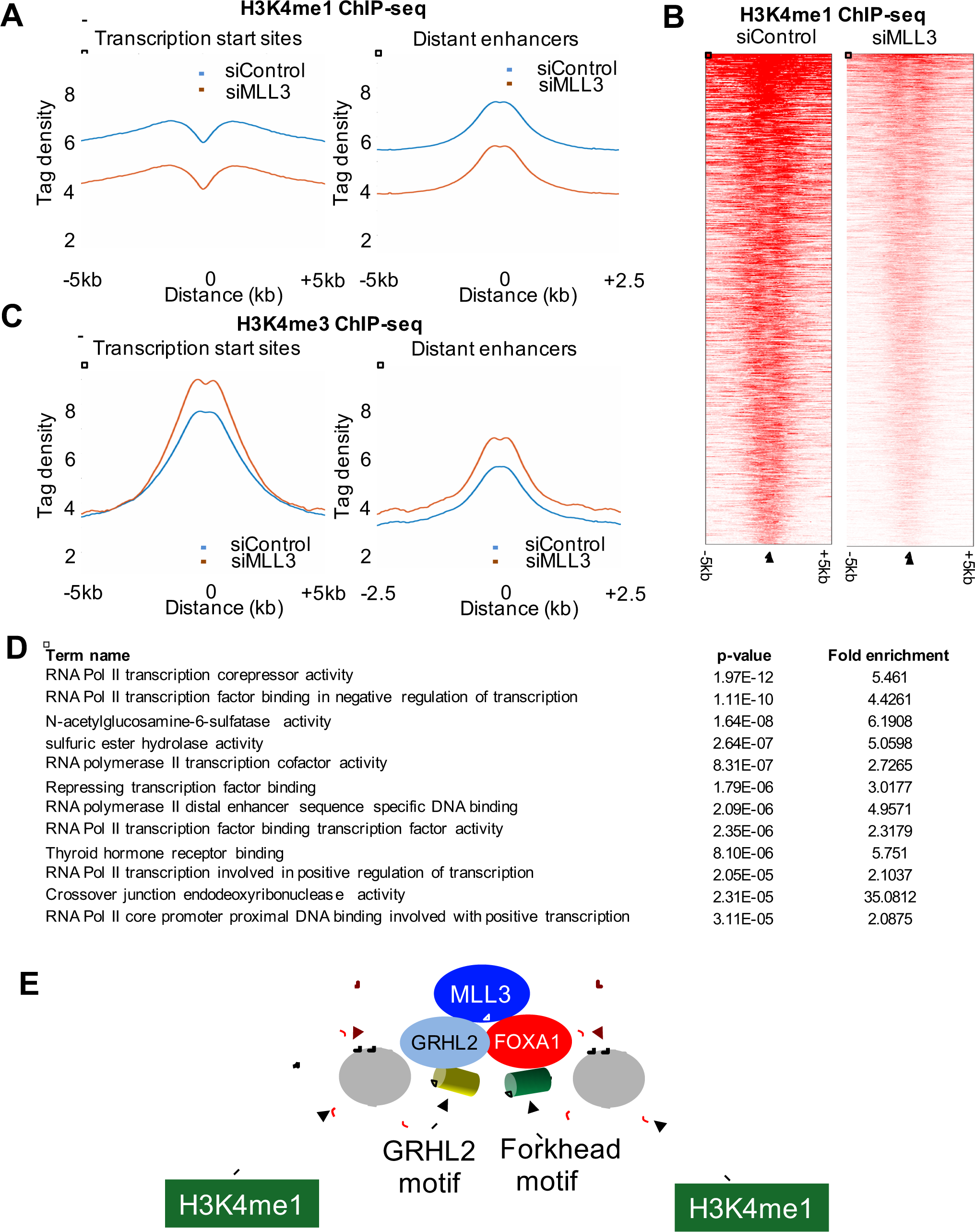
Direct dependency of H3K4me1 on MLL3 at enhancers. The effect of MLL3 silencing on H3K4me1 and H3K4me3 binding were assessed by ChIP-seq in siControl or siMLL3 transfected cells. Differential H3K4me1 and H3K4me3 peaks that were altered by silencing of MLL3 were identified. **A.** The average signal intensity of H3K4me1 at enhancer elements or promoters following silencing of MLL3. **B.** Heatmap of differential H3K4me1 regions that occur at FOXA1/MLL3 co-bound regions. **C.** The average signal intensity of H3K4me3 at enhancer elements or promoters following silencing of MLL3. **D.** Enriched pathways within the 776 FOXA1/MLL3 co-bound reigons that had decreased H3K4me1 signal following silencing of MLL3. **E.** Model showing FOXA1 and GRHL2 recruitment of MLL3, which subsequently contributes to monomethylation of H3K4.

## DISCUSSION

We propose a mechanism whereby the pioneer factor FOXA1 interacts with chromatin and recruits the methyltransferase MLL3, faciliating the deposition of H3K4me1 in breast cancer cancer cells (Figure 4E). This is a new mechanism where an enhancer-specific pioneer factor FOXA1 can interact with a chromatin modifier (MLL3) to facilitate the occurrence of H3K4me1 histone mark at regions that become functional enhancer elements. We have described a novel link between FOXA1, MLL3 and H3K4methylation, revealed by RIME, an unbiased proteomic technique, that showed MLL3 to be a robust FOXA1 interacting protein in MCF-7 breast cancer cells. MLL3 has been shown to contribute to H3K4me3 at promoter regions (Ananthanarayanan et al., 2011), but our evidence would suggest that MLL3 can also contribute to H3K4me1 marks at enhancer regions and that this is determined by the transcription factors that recruit MLL3 to the chromatin. Two mechanisms for MLL3 recruitment to chromatin are revealed by mining of MLL3 ChIP-seq data. Motifs for Forkhead and GRHL2 transcription factors were identified. Interestingly, MLL3-chromatin occupany was shown to occur via FOXA1, GRHL2 or both factors and our functional experiments confirm that FOXA1 is essential for MLL3 binding. GRHL2 has been implicated in metastasis in breast cancer (Werner et al., 2013; Xiang et al., 2012) and we hypothesize that its influence on cell migration and metastatic potential is attributed to its ability to tether MLL3 at chromatin and mediate enhancer activity. Whether GRHL2 is involved in ER/FOXA1+ breast cancer, or can function independently of FOXA1 (and ER) is not clear, but GRHL2 is located on chromosome 8 and is commonly co-amplified with c-Myc suggesting that any role for GRHL2 in mediating recruitment of the enzyme MLL3 may be substantially altered in tumors with the commonly occuring chromosome 8 amplification.

The predominant paradigm is that H3K4me1 and H3K4me2 marks are signatures of enhancer regions, whereas H3K4me3 is enriched at the promoters of coding genes (Calo and Wysocka, 2013; Heintzman et al., 2009, 2007). Our findings would suggest that the presence of H3K4me1 marks at enhancers are enriched at FOXA1 bound enhancer elements because this pioneer factor is able to recruit the enzyme that contributes to the deposition of this methylation event biased towards regions co-bound by FOXA1, GRHL2 and the methyltransferase MLL3. Recently, it has been reported that MLL3/4 contributes to monomethylation (H3K4me1) of promoter regions in myoblasts (Cheng et al., 2014). It has also been shown that Trr, the Drosophila homolog of the mammalian MLL3/4 COMPASS-like complexes, can function as a major H3K4 monomethyltransferase on enhancers in vivo (Herz et al., 2012), with a modest decrease of H3K4me1 in mouse embryonic fibroblasts (MEFs) from MLL3 knockout mice (Herz et al., 2012). In our breast cancer cells, we see a pronounced depletion of H3K4me1 following MLL3 silencing. Since MLL3 and the related protein MLL4 function as a complex, it is possible that both MLL3 and MLL4 contribute to the enhancer H3K4 methylation marks, as both proteins needed to be deleted in MEFs to see the decrease in the H3K4me1 (Herz et al., 2012). However, we did not find MLL4 as a FOXA1 interacting protein and even in MLL3 depleted cells, there was no FOXA1-MLL4 interactions (data not shown) suggesting a lack of redundancy between MLL3 and MLL4 in our breast cancer cells. It has also been shown that unlike MLL3, the depletion of MLL4 had no effect on the estrogen-mediated activation of HOXC6 (Ansari et al., 2011), suggesting that they are not functionally linked in ER biology. The specific role for MLL3 in ER+ breast cancer is supported by the recent finding that MLL3 was mutated in 5 out of 46 ER-positive breast cancer samples (Ellis et al., 2012) and although the mutations occur at distinct regions within MLL3, a common result is putative pertubation in key enzymatic domains within MLL3. The functional significance of these mutations warrants investigation, although the large size of MLL3 (541kDa) makes these functional experiments a challenging endeavour.

Taken together, we propose a new mechanism, where the pioneer factor FOXA1 interacts with the chromatin modifier MLL3 to facilitate monomethylation of H3K4 at enhancer elements, resulting in the potential for transcription from these enhancer regions. These findings imply that the transription factors that associate with enhancer elements are capable of actively contributing to the H3K4me1 that occurs at enhancers, rather than requiring their presence for chromatin occupancy.

## METHODS

### Cell lines

MCF-7 cells were obtained from ATCC. MCF-7 were grown in Dulbecco’s Modified Eagle Medium (DMEM) supplemented with 10% heat-inactivated foetal bovine serum (FBS), 2 mM L-glutamine, 50U/ml penicillin and 50μg/ml. All cell lines were regularly genotyped using STR profilling (Promega GenePrint 10 system). Cell lines were regularly tested for mycoplasma infection.

### Antibodies

The antibodies used for ChIP-seq were anti-FOXA1 (ab5089) from Abcam, anti-H3K4me1 (05-1248-S) from Millipore, anti-H3K4me3 (05-1339) from Millipore and anti-GRHL2 (HPA004820) from Sigma Aldrich. The custom anti-MLL3 antibody was provided by Prof. Ali Shilatifard (Stowers Institute, Kansas City, USA and Northwestern University Feinberg School of Medicine, Chicago, USA).

### Chromatin immunoprecipitation sequencing (ChIP-seq)

ChIPs were performed as described previously (Schmidt et al., 2009). The ChIP-seq and the input libraries were prepared using the TruSeq ChIP Sample Prep Kit (Illumina). ChIP-seq of each factor was performed in at least biological triplicates. Reads were mapped to hg19 genome using Gsnap version 2015-09-29 (Wu TD1, Nacu S., 2010). Aligned reads with the mapping quality less than 5 were filtered out. The read alignments from three replicates were combined into a single library and peaks were called with MACS2 version 2.0.10.20131216 (Zhang Y1 at al., 2008) using sequences from MCF7 chromatin extracts as a background input control. For the ChIP samples of Histones with mono and tri-methyl modifications the broad peaks were called. The peaks yielded with MACS2 q value <= 1e-5 were selected for downstream analysis. Meme version 4.9.1 (Timothy et al., 2009) was used to detect known and discover novel binding motifs amongst tag-enriched sequences. For visualizing tag density and signal distribution heatmap the normalized tumour read coverage in a window of +/− 2.5 or 5 kb region flanking the tag midpoint was generated using the bin size of 1/100 of the window length. Differential binding analysis (Diffbind) was performed as described previously (Brown and Stark, 2011).

For ChIP-qPCR, primer sequences used were: TFF1 Fwd: GTGGTTCACTCCCCTGTGTC, TFF1 Rev: GAGGCATGGTACAGGAGAGC, GREB1 Fwd: CACGTCCCCACCTCACTG, GREB1 Rev: TGTTCAGCTTCGGGACACC, PGR Fwd: GCTCCAGCTAACTGATGGTCTG, PGR Rev: TGGGCCTAGATTATTGAGTTCAGG.

### Rapid IP Mass-spectrometry of Endogenous proteins (RIME)

RIME was performed as previously described (Mohammed et al., 2013). Proteins were digested using Trypsin. Maximum allowed missed cleavage was 2, the peptide threshold was 95% and the protein False Discovery Rate (FDR) was set to 0.5%. Proteins were considered as interactors when at least 2 high-confident peptides were identified and when none of these peptides were observed in matched IgG control RIME experiments. Additionally, FOXA1 interactors were filtered using the CRAPOME database (www.crapome.org).

### Small interfering RNA transfections

Small interfering RNA (siRNA) used to silence FOXA1 were obtained from Dharmacon RNAi Technologies. The sequence of the siRNA that targeted FoxA1 is GAGAGAAAAAAUCAACAGC and has been previously validated (Hurtado et al., 2011). Small interfering Smartpool RNA used to silence MLL3 were obtained from Thermo Scientific (L-007039-00-0020). AllStars Negative Control siRNA (Qiagen) was used as a negative controls. Cells were transfected with siRNA using Lipofectamine 2000 (Invitrogen).

### Preparation of mRNA

Cells cultured in 15 cm^2^ dishes were first washed twice with cold PBS and RNA was extracted using RNeasy kit (Qiagen) as per manufacturer’s instructions. DNA was degraded by adding 20 units of RNAse free DNAseI (Roche Diagnostics GmbH) for 15 minutes at room temperature. DNAse I treatment was performed on columns.

### Preparation of cDNA

1μ of total RNA was diluted to a final volume of 11 μl using 100 μ of random primers (Promega), 2.5mM of dNTP mix and nuclease-free water. This mixture was then incubated at 65°C for 5 minutes. First strand buffer (Invitrogen) and 10mM 1.4-dithiothreitol (DTT) (Invitrogen) was then added to this mixture and incubated at 25°C for 10 minutes to allow primer annealing. This mixture was then heated at 42°C for 1 minute and 200 units of SuperScriptTM III Reverse Transcriptase (Invitrogen) were added. This final mixture was then incubated at 42°C for an additional 50 minutes and the process was stopped after inactivating the enzyme at 70°C for 15 minutes. The resulting cDNA was then diluted 1:10 in H_2_O for subsequent use.

### Quantitative RT-PCR

Quantitative PCR was performed using a Stratagene^®^ Mx3005P RealTime machine. Each qPCR reaction contained Power SYBR^®^ Green PCR Master Mix (Applied Biosystems, California, USA), 250nM of each primer, 2μl of DNA eluted after chromatin IP and nuclease-free H2O added to a final volume of 20μl. The PCR program consisted of first heat activating the Taq polymerase at 95°C for 10 minutes. This was then followed by 45 cycles of 15 seconds at 95°C and 30 seconds at 60°C. The fluorescence from each well was analysed at every cycle. The final step involved increasing the temperature from 65°C to 95°C and continuously reading the fluorescence. Reactions were performed in triplicates and results were analysed using the delta-delta Ct method (Livak and Schmittgen, 2001).

## Acknowledgements

The authors would like to thank the members of the proteomic, genomics and bioinformatic core facilities at Cancer Research UK for their help. We would like to thank Prof Ali Shilatifard (Northwestern University) for generously providing MLL3 antibody. We would like to thank Aisling Redmond for critically reading the manuscript. The authors thank Gordon Brown for assistance with ChIP-seq and proteomics data analysis. We would like to acknowledge the support of the University of Cambridge, Cancer Research UK and Hutchison Whampoa Limited. K.M.J is funded by Cancer Research UK. J.S.C. is supported by an ERC Consolidator grant and an EMBO Young investigator award.

## Data deposition

All ChIP-seq data is deposited in GEO with accession number to be provided.

All proteomic data is deposited with the PRIDE database with accession number to be provided.

## Author contributions

K.M.J. and J.S.C designed experiments. K.M.J., S.N. and A.A.S performed experiments. K.M.J., I.C and S.N. analyzed the data. K.M.J. and J.S.C wrote the manuscript, with input from all authors. J.S.C oversaw the work.

